# Perturbations in Risk/Reward Decision Making and Frontal Cortical Catecholamine Regulation Induced by Mild Traumatic Brain Injury

**DOI:** 10.1101/2024.02.22.581581

**Authors:** Christopher P. Knapp, Eleni Papadopoulos, Jessica A. Loweth, Ramesh Raghupathi, Stan B. Floresco, Barry D. Waterhouse, Rachel L. Navarra

**Author notes:** **Contact Information (full mailing address, and e-mail):** Christopher P. Knapp; 42 East Laurel Road, Suite 2200, Stratford, NJ 08084; Eleni Papadopoulos; 42 East Laurel Road, Suite 2200, Stratford, NJ 08084; Jessica A. Loweth; 42 East Laurel Road, Suite 2200, Stratford, NJ 08084; Ramesh Raghupathi; 2900 W. Queen Lane, Philadelphia, PA 19129; Stan B. Floresco; 2136 West Mall, Vancouver BC, V6T 1Z4, Canada; Barry D. Waterhouse; 42 East Laurel Road, Suite 2200, Stratford, NJ 08084; Rachel L. Navarra; 42 East Laurel Road, Suite 2200, Stratford, NJ 08084. **Corresponding author: Christopher Knapp (contact)**. **Co-Corresponding author: Rachel Navarra (contact)**.

## Abstract

Mild traumatic brain injury (mTBI) can disrupt cognitive processes that influence risk taking behavior. Athletes, military personnel, and domestic violence victims often experience multiple mTBIs; however, little is known regarding the effects of repetitive injury (rmTBI) on risk/reward decision making or whether these outcomes are sex specific. Risk/reward decision making is mediated by the prefrontal cortex (PFC), which is composed of several sub-regions including the medial PFC (mPFC), anterior cingulate cortex (ACC), and orbitofrontal cortex (OFC). These regions are densely innervated by catecholaminergic fibers, which modulate PFC-mediated cognitive processes. Aberrant catecholamine activity within the PFC has been documented following TBI, which may underlie TBI-induced risky behavior. Tyrosine hydroxylase (TH) and norepinephrine transporter (NET) regulate catecholamine homeostasis within the PFC; however, it has not been determined how rmTBI affects these proteins. The present study aimed to characterize the effects of rmTBI on risk/reward decision making behavior and catecholamine transmitter regulatory proteins within the PFC. Risk/reward decision making was evaluated using a probabilistic discounting task (PDT) which required rats to choose between small/certain rewards delivered with 100% certainty and large/risky rewards delivered with decreasing probabilities over a session. Rats were first trained on the PDT and then exposed to sham, single (smTBI), or a series of three closed-head control cortical impact (CH-CCI) injuries over the course of one week, followed by four weeks of PDT testing. In week 1 post-final surgery, mTBI generally enhanced preference for the larger/riskier option with these effects seemingly more prominent in females. These effects resolved by week 2 post-final surgery indicating that the effects of mTBI on choice behavior are transient. By week 4, males, but not females, exhibited increased latencies to make riskier choices following rmTBI, demonstrating a delayed effect of injury on information processing speed. A separate group of rats was used to measure changes in levels of TH and NET within the mPFC, ACC, and OFC forty-eight hours after mTBI. No injury-induced differences were observed within the mPFC or ACC. In the OFC, females exhibited dramatic increases in TH levels following smTBI, but only small increases following rmTBI. Both males and females; however, experienced reduced levels of NET following rmTBI, which may function as a compensatory response to increased extracellular levels of catecholamines. Together, these results suggest that OFC is more susceptible to catecholamine instability after rmTBI, a finding indicating that not all areas of the PFC contribute equally to the observed TBI-induced catecholamine imbalances. Overall, combining the CH-CCI model of rmTBI with the PDT proved effective in revealing time-dependent and sex-specific changes in risk/reward decision making and catecholamine regulation following repetitive mild head injuries.

## Introduction

Athletes, military personnel, and domestic violence victims often experience multiple traumatic brain injuries (TBIs)^1–11^, the majority of which are classified as mild (mTBIs), or concussions^3–6,10,12^. While the effects of single injuries are often transient, evidence suggests that repetitive mild injuries (rmTBI) can result in more severe and longer-lasting cognitive impairments^5,9,13–15^. The prefrontal cortex (PFC), which is most frequently affected following TBI events^16,17^, mediates complex cognitive processes that regulate decision making and action in situations that involve uncertain risk/reward outcomes. The Iowa Gambling Task (IGT)^18^ has primarily been used in clinical assessments of risk/reward decision making in TBI patients. This task models real-life decisions involving reward, punishment, and uncertainty of outcomes. In this task, participants choose cards from four different decks in an opportunity to win money. Two decks are associated with large gains but also pose risks of losses that are more frequent or of higher magnitude, and two decks are associated with small gains, but pose less risk of loss. During task performance, healthy individuals learn that choosing the small/certain decks are advantageous to maximize overall gain net gain. Initial studies established that patients who have experienced TBI, especially with damage encroaching on the PFC, displayed suboptimal decision profiles, resulting in greater loss of money on the IGT^18–20^. Further clinical studies have consistently reported increased risk-taking behavior in single TBI cases^18–23^; although the effects of repetitive mild injuries have not been explored. One limitation of clincial studies on how cognitive functions may be affected by repetitive TBI it that it is difficult to control for the severity, timing, and number of injuries that a TBI patient suffers. Moreover, much of the current TBI research does not include a proportionate number of men and women in their assessments, despite numerous reports of sex differences in TBI susceptibility rates^24–37^, post-injury symptoms^27,28,38–40^, and recovery patterns^28,38–42^. Post-TBI assessments of decision making often group men and women together^18–20,22,23^, making it difficult to differentiate potential sex differences in risk/reward decision making following injury. These deficiencies underscore the need for the development of pre-clinical models that can appropriately access, reveal, and differentiate potential sex-specific deficits in PFC-mediated cognitive processes following TBI.

Clinical and pre-clinical investigations have revealed that the PFC contains multiple interconnected regions that work together to facilitate efficient risk/reward decisions^43–57^. These areas include anterior cingulate cortex (ACC), regions of the medial PFC (mPFC), and orbitofrontal cortex (OFC). Individual damage to any of these regions has been shown to impair reward-guided learning and action selection^44,53^, increase perseverative behavior^18,58^, reduce sensitivity to consequences of risky decisions^18,20^, and impair information processing relating to new and/or changing risk/reward contingencies^22,23,55^. TBI often damages multiple areas of the PFC, which causes collective impairments to decisional processes, leading to the reported increases in risk taking behavior. While the effects of repetitive TBI have not yet been explored, it is likely that multiple head injuries exacerbate existing damage to prefrontal areas and cause new injuries in previously uninjured regions.

The PFC receives dense innervation from catecholaminergic fibers, containing dopamine (DA) and norepinephrine (NE), which modulate PFC-mediated processes^59–64^. An optimal balance of catecholaminergic signaling is required for normal operation of all PFC regions. While small increases in DA and NE neurotransmission can improve cognitive performance^65–70^, catecholamine activity outside this optimal range can result in impaired cognitive functioning. Previous reports have demonstrated aberrant catecholamine activity within the prefrontal cortex following TBI^71,72^, suggesting that imbalanced DA and NE levels may underlie injury-induced risky decision making. The reasons for these alterations are unclear; however, some studies have reported changes in catecholamine-associated regulatory protein levels following TBI. Increased levels of tyrosine hydroxylase (TH), the rate-limiting enzyme for catecholamine synthesis, have been observed throughout the brain^71,73–76^, suggesting a surge in catecholamine production. Reuptake transporter proteins, DAT and NET, remove extracellular DA and NE after their release to maintain efficient catecholaminergic signaling. NET, in particular, is the primary transporter for DA and NE in the PFC^77–79^; however, no studies have examined changes in NET expression after TBI despite brain-wide, and sometimes sex-dependent, decreases in DAT being reported^74,80–82^. Furthermore, how repetitive injury affects these proteins and whether changes in TH and NET expression are similar across all prefrontal areas remains to be explored. Given the dissociable contributions of the mPFC, ACC, and OFC to risk/reward decision making, differential disruptions to catecholamine activity within each sub-region would result in distinct changes to the processing and execution of probabilistic-based decisions.

The goal of the present study was to characterize the effects of rmTBI on risk/reward decision making and to identify potential neural mechanisms of catecholamine transmitter imbalances in the PFC underlying these injury-induced effects. To this end, we utilized an established pre-clinical assay of risk/reward decision making, i.e. the probabilistic discounting task (PDT)^83^. This task requires rats to choose between small/certain rewards delivered with 100% certainty and large/risky rewards delivered with probabilities that decrease across a session. Like the IGT, performance on the PDT relies on intact PFC functioning, and changes in neural^55,57^ or catecholaminergic^45,56^ activity within the PFC have been shown to alter these types of risk-related decisions. This sensitivity to changes in PFC activity makes the PDT a viable assay for detecting potential impairments to risk/reward decision making following TBI. Western blotting procedures were then used to determine levels of catecholamine-associated regulatory proteins within specific subregions of the PFC known to influence decision making processes at the time point of first behavioral evaluation following injury. Together, these evaluations established a framework for investigating sex-specific changes in PFC-mediated executive function and catecholamine regulation following repetitive mild head injuries.

## Methods

### Animals

Fifty-five male and fifty-four female Long-Evans rats were used in this study. Animals were obtained at either 3-4 weeks old/50-75g (behavioral experiments) and 5-6 weeks old/100-125g (Western blotting experiments) from Charles River Laboratories and housed in a 12h:12h reverse light/dark cycle facility. Following a week of acclimation, rats were single housed into separate cages, and placed on a food regulated diet (5g per 100g body weight/day) with *ad libitum* access to water. They were maintained to 85% of their free feeding weight throughout the duration of these studies. All experimental procedures were in accordance with the Rowan University School of Osteopathic Medicine Institutional Animal Care and Use Committee and the National Institutes of Health Guide for the Care and Use of Laboratory Animals.

### Probabilistic Discounting Task (PDT)

#### Apparatus

Behavioral studies were conducted in 16 operant chambers [29cm (L) x 24cm (W) x 29cm (H); Med-Associates, Albans, VT] enclosed within sound attenuating boxes. Operant chambers were equipped with a fan, a house light, and 2 retractable levers located on either side of a food dispenser where sucrose pellet rewards (45 mg; Bio-Serv, Flemington, NJ) were delivered. A photo beam was located at the dispenser entry point to detect reward collection. Custom built Med Associates Nexlink computer packages controlled the training, testing, and data acquisition during performance.

#### Lever-Pressing Training

Initial training protocols were adapted from St. Onge and Floresco^83^. Age-matched male (n = 31) and female (n = 30) rats were first trained to press a single lever (either the left or right) using a fixed-ratio one (FR1) schedule to a criterion of 50 presses within 30 minutes. Once criterion was achieved, rats repeated this procedure for the opposite lever. Rats then trained on a simplified version of the PDT (90 trials per session) which required them to press one of the two levers within a 10 second period for a sucrose reward delivered with a 50% probability. This procedure familiarized them with the probabilistic nature of actions and outcomes. Rats were trained for at least 3 days to a criterion of 75 or more successful trials (i.e.; ≤ 15 omissions) on the simplified PDT.

#### PDT Training and Testing

The PDT was used to assess changes in risk/reward decision making and has been described previously^55–57,83–87^ (Fig. 1). This task required rats to choose between levers that result in either small/certain rewards (1 pellet) delivered with 100% certainty and large/risky rewards (4 pellets) delivered with decreasing probabilities across a series of five trial blocks (i.e. 100% probability → 50% → 25% → 12.5% → 6.25%). Each session took 52.5 minutes to complete and consisted of 90 trials, separated into 5 blocks of 18 trials. These 18 trials consisted of 8 forced-choice trials where only one lever was extended allowing rats to learn the relative likelihood of obtaining the larger reward in each block. This was followed by 10 free-choice trials, where both levers were extended allowing rats to freely choose between the small/certain or the large/risky lever. Each session began in darkness with both levers retracted. A trial began every 35 seconds with the illumination of the house light and extension of one or both levers. Once a lever was chosen, both levers retracted, rats were rewarded 1 pellet if they chose the small/certain lever or a possible 4 pellets if they chose the large/risky lever, and the house lights turned off. If the rat did not respond within a 10 second period, levers retracted, and the house light turned off until the next trial and the trial was scored as an omission.

**Figure 1.**
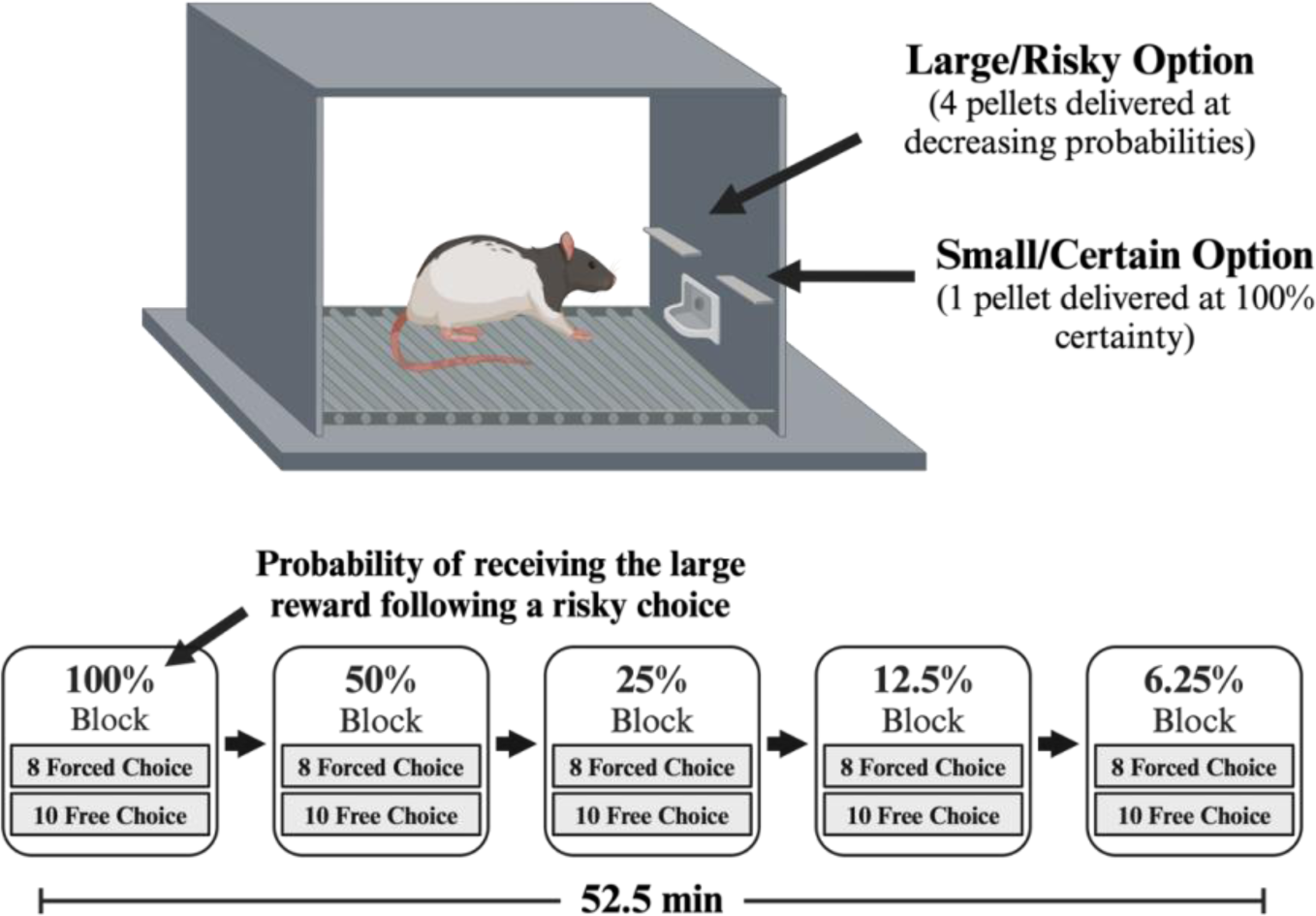
Schematic of the Probabilistic Discounting Task. **Top:** Cost/benefit contingencies associated with each lever. **Bottom:** Task design depicting the decreasing large/risky reward probabilities across five trial blocks.

Rats were trained 7 days per week until each cohort achieved baseline criteria, which included choosing the risky lever in >80% of trials in the 100% block and maintaining stable patterns of choice for 3 consecutive sessions. Determining stable baseline performance involved analyzing 3 consecutive sessions using a repeated measures analysis of variance (ANOVA) with two within-subjects factors (day and trial block). Rats were required to demonstrate a significant main effect of block (p < 0.05), but not a main effect of day nor a day x block interaction (p > 0.1). If animals, as a group, met these 3 requirements, they were determined to have achieved stable baseline levels of choice behavior.

Rats required ∼17 days of training before stable criterion performance was achieved. They then received TBI or sham surgical procedures. In so doing, animals were rank ordered based on the average ratio of large/risky lever presses over successful free-choice trials across the last three training days. Animals with the highest ratio were assigned a higher rank. A reverse Latin square method was then used to assign rats to one of three surgical groups: sham (uninjured), single injury (smTBI), or repetitive injury (rmTBI). This method was used to limit bias when forming surgical groups by having equal amounts of risky and risk-adverse rats in each group. Forty-eight hours after the final surgery, animals were reintroduced to the PDT and tested 5 days per week for four weeks to assess changes in risk/reward decision making.

### Surgery

On the day of surgery, each group of rats (9-10 weeks old) underwent a total of three surgeries over the course of one week separated by 2 days using the closed head-controlled cortical impact (CH-CCI) model (Custom Design & Fabrication Incorporated, Glenn Allen, VA). Briefly, animals were anesthetized at 4% isoflurane in 95% oxygen/5% carbon dioxide, then maintained at ∼2.0% isoflurane while a midline incision of 2cm was made to expose the skull. Anesthesia was discontinued and the rat was transferred to a stage under the CCI device with its head resting on a foam pad [10cm (L) x 5cm (W) x 1 cm (H)]. A 5mm diameter metal impactor tip was positioned on the skull surface along the sagittal suture with the edge of the tip aligned 2 mm behind bregma. The tip was then electronically driven at a velocity of 5.5m/s and a depth of 2.5mm below the surface point of contact with a dwell time of 100ms. Following injury, righting reflex times were recorded by measuring the latency for animals to regain normal posture after being placed in the supine position. Animals were then re-anesthetized and the incision was closed with wound clips. rmTBI rats received an impact on all 3 surgery days whereas smTBI rats received sham surgeries on the first 2 surgery days and an impact on the 3^rd^ day. Sham rats underwent the same surgical procedures but were not impacted on any day.

### Western Blotting

To examine the effects of rmTBI on TH and NET levels within sub-regions of the PFC, separate groups of age-matched male (n = 24) and female (n = 24) rats were subjected to the same surgical procedures as described above. Forty-eight hours after their final surgery, anesthetized animals were decapitated and the mPFC, ACC, and OFC were dissected^88^ (Fig. 2), immediately frozen on dry ice, and stored at -80°C until use. Tissue was then homogenized in lysis buffer (consisting of 10mM HEPES, 2mM EDTA, 2mM EGTA, 1% Triton X-100, 1X Protease Inhibitor I, and 1X Protease Inhibitor II), followed by refrigerated centrifugation at 20,000 x g for 2 minutes at 4°C to generate whole cell lysates^89–91^. The supernatants were then aliquoted and stored at -80°C until use. Protein concentrations were determined using a Bio-Rad DC Protein Assay Kit (Bio-Rad, Hercules, CA). Protein samples were prepared in 4X sample buffer (Bio-Rad, Hercules, CA) and heated at 90° for 3 minutes. Equal amounts of protein (15ug) were loaded into each well of a Criterion XT Bis-Tris Protein Gel (Bio-Rad, Hercules, CA). Following gel electrophoresis, protein were transferred to Immuno-Blot PVDF membranes (Bio-Rad, Hercules, CA). Membranes were blocked with 5% non-fat dry milk in TBS-T (20X TBS and 10% Tween) for 1 hour and then probed with either rabbit anti-TH (1:1000; MilliporeSigma, Temecula, CA) or rabbit anti-NET (1:1000; Abcam, Waltham, MA) antibody at 4°C overnight followed by goat anti-rabbit secondary antibody conjugated with peroxidase (1:10,000; Rockland Immunochemicals, Inc., Limerick, PA) for 1 hour the next day. β-actin (1:2000; MilliporeSigma, Temecula, CA) was used as the loading control. Chemiluminescence was detected using Clarity Western ECL substrate (Bio-Rad, Hercules, CA), imaged using Azure c400 Biosystems imaging system (Azure Biosystems, Dublin, CA), and analyzed using AzureSpot Analysis Software (Azure Biosystems, Dublin, CA).

**Figure 2.**
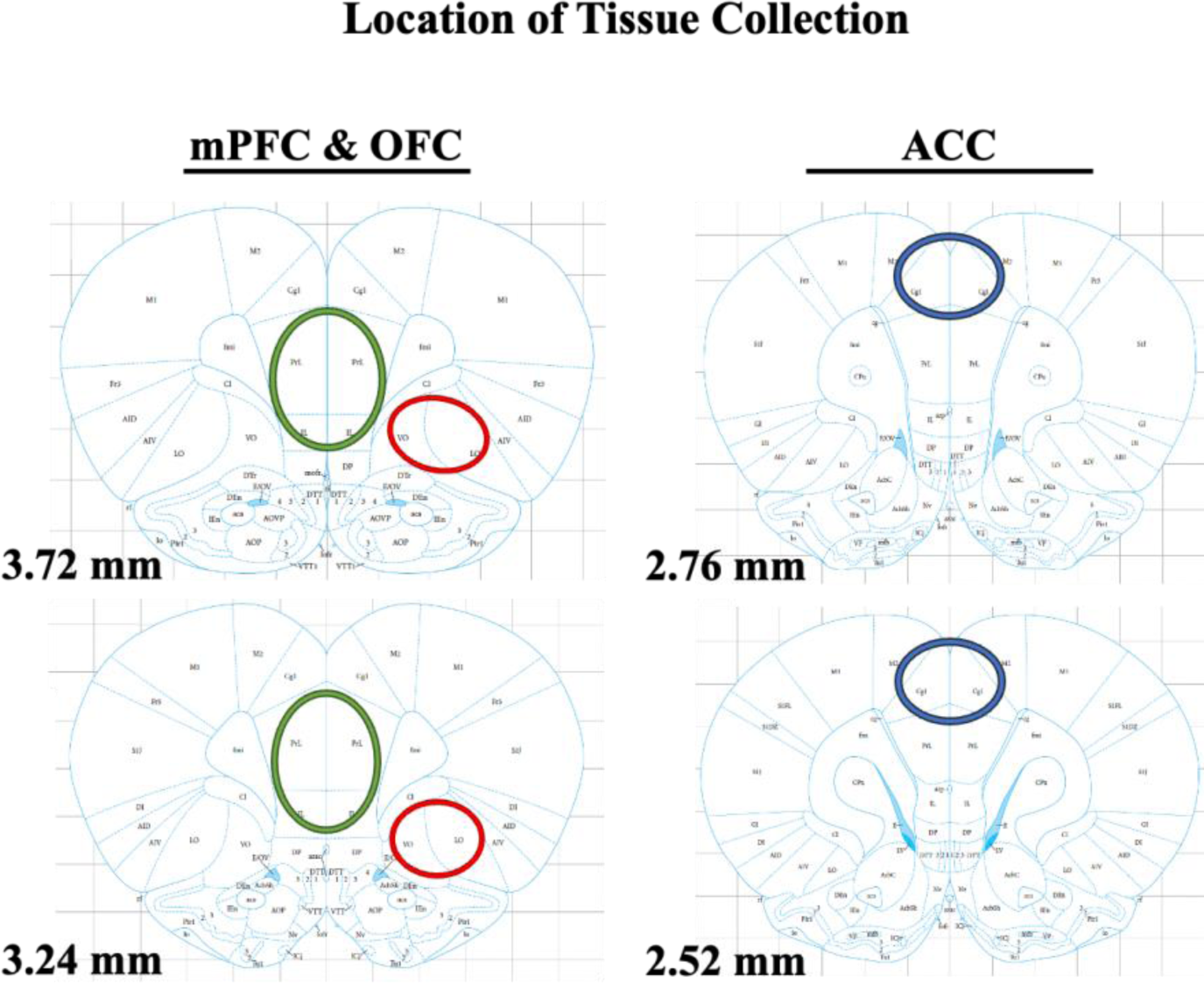
Location of tissue collected from the medial prefrontal, orbitofrontal, and anterior cingulate regions for Western Blot assays (adapted from Paxinos and Watson, 2007).

### Statistical analysis

Analysis of righting reflex and Western blotting data were performed using GraphPad Prism software (GraphPad Software, San Diego CA). Analysis of choice behavior and choice latencies were performed using SPSS software (IBM, SPSS Inc.). For all surgery groups, righting reflex times, measured as the latency to regain sternal recumbency following impact, were averaged together, respectively, within each surgery day. Male and female righting reflex data were then analyzed separately using two-way repeated measures ANOVAs with surgery day (day 1, day 2, and day 3) as the within-subjects factor and injury condition (sham, smTBI, and rmTBI) as the between-subjects factor. For post-surgery PDT behavior, choice behavior was measured as the percentage of choices directed towards the large/risky option, while choice latency was measured as the latency to press either the large/risky or small/certain lever. These data were averaged across 3 consecutive sessions, respectively, and analyzed using three-way mixed-design ANOVAs with trial block (100%, 50%, 25%, 12.5%, and 6.25%) as the within-subjects factor and injury condition (sham, smTBI, and rmTBI) and sex (male and female) as the between-subjects factor. The effect of trial block was always significant (p < 0.05) and will not be discussed further. For Western blotting, individual one-way ANOVAs were used to analyze group differences (sham vs. smTBI vs. rmTBI) in protein levels for each PFC sub-region. This analysis was performed for males and females. Dunnett’s multiple comparisons tests, when appropriate, were used to compare individual differences when overall significance was found. For all results, statistical significance was determined by a p value < 0.05.

## Results

### Acute Response to Injury

Immediately following sham injury or mTBI, the latency to regain righting reflex was recorded. rmTBI in male rats resulted in longer righting reflex times compared to sham animals across all three surgical days (Table 1). Analysis of righting reflex data revealed significant main effects of both day [F (1.635, 85.02) = 10.11, p = 0.0003] and injury [F (2, 52) = 5.892, p = 0.0049] as well as a significant day x injury interaction [F (4, 104) = 4.901, p = 0.0012]. Dunnett’s multiple comparisons analysis revealed that rmTBI rats demonstrated longer righting reflex times compared to sham rats on days 1 (p = 0.0228) and 2 (p = 0.0538) whereas smTBI rats displayed longer righting reflex times on day 3 (p = 0.0007).

**Table 1.** Average righting reflex times (seconds) of male and female sham, single (smTBI), and repetitive (rmTBI) injury groups across all three surgery days. On surgery day 1, repetitive injured (rmTBI) males and females exhibited longer righting reflex times compared to their respective sham groups. On surgery day 2, rmTBI males demonstrated a trend towards longer righting reflex times, whereas rmTBI females demonstrated a significant increase in righting reflex times. On surgery day 3, smTBI males and females demonstrated longer righting reflex times compared to their respective sham groups. Values represent mean ± SEM. * denotes p < 0.05 and ^ denotes p < 0.1 from shams analyzed with Dunnet’s multiple comparisons tests.

| Sex | Injury Condition | N | Righting Reflex (sec) |  |  |
| --- | --- | --- | --- | --- | --- |
|  |  |  | Surgery Day 1 | Surgery Day 2 | Surgery Day 3 |
| Male | Sham | 19 | 350.5 ± 48.4 | 233.8 ± 25.0 | 245.0 ± 18.1 |
| Female | Sham | 19 | 344.2 ± 39.5 | 237.0 ± 28.9 | 207.7 ± 22.5 |
| Male | smTBI | 18 | 328.8 ± 31.4 | 315.1 ± 35.4 | 397.5 ± 32.7* |
| Female | smTBI | 17 | 255.0 ± 31.0 | 191.2 ± 21.8 | 320.7 ± 21.9* |
| Male | rmTBI | 18 | 507.3 ± 33.6* | 321.9 ± 29.5^ | 326.2 ± 35.5 |
| Female | rmTBI | 18 | 500.6 ± 32.9* | 367.4 ± 34.3* | 234.3 ± 28.3 |

mTBI in female rats also resulted in longer righting reflex times compared to sham animals across all three surgical days (Table 1). Analysis of righting reflex data revealed significant main effects of both day [F (1.962, 100.1) = 16.74, p < 0.0001] and injury [F (2, 51) = 9.049, p = 0.0004] as well as a significant day x injury interaction [F (4, 102) = 10.98, p < 0.0001]. Dunnett’s multiple comparisons analysis revealed rmTBI rats demonstrated longer righting reflex times compared to sham rats on days 1 (p = 0.0086) and 2 (p = 0.0121) whereas smTBI rats displayed longer righting reflex times on day 3 (p = 0.0019).

### Effects of mTBI on Choice Behavior

In the first week post-final surgery (Fig. 3), analysis of choice data revealed a significant main effect of injury [F (2, 55) = 3.582, p = 0.034] with no injury x block, or three-way interaction with the sex factors (all ps > 0.05). There were also no main effects of sex [F (1, 55) = 2.424, p = 0.125] or injury x sex interaction [F (2, 55) = 0.303, p = 0.740]. The main effect of injury reflected the observation that TBI increased risky choice during Week 1 of testing relative to the sham group. This was confirmed with Dunnett’s multiple comparisons analysis, that revealed smTBI rats displayed a significant increase risky choice (p = 0.035) in comparison to sham rats, and a trend towards a significant increase in risky choice following rmTBI (p = 0.073). Although we did not observe significant interactions with the sex factor, visual inspection of the data plotted separately for each sex (Fig. 3A, top left inset) suggest that the increase in risky choice was more robustly in the female injury groups in the first week post-final injury.

**Figure 3.**
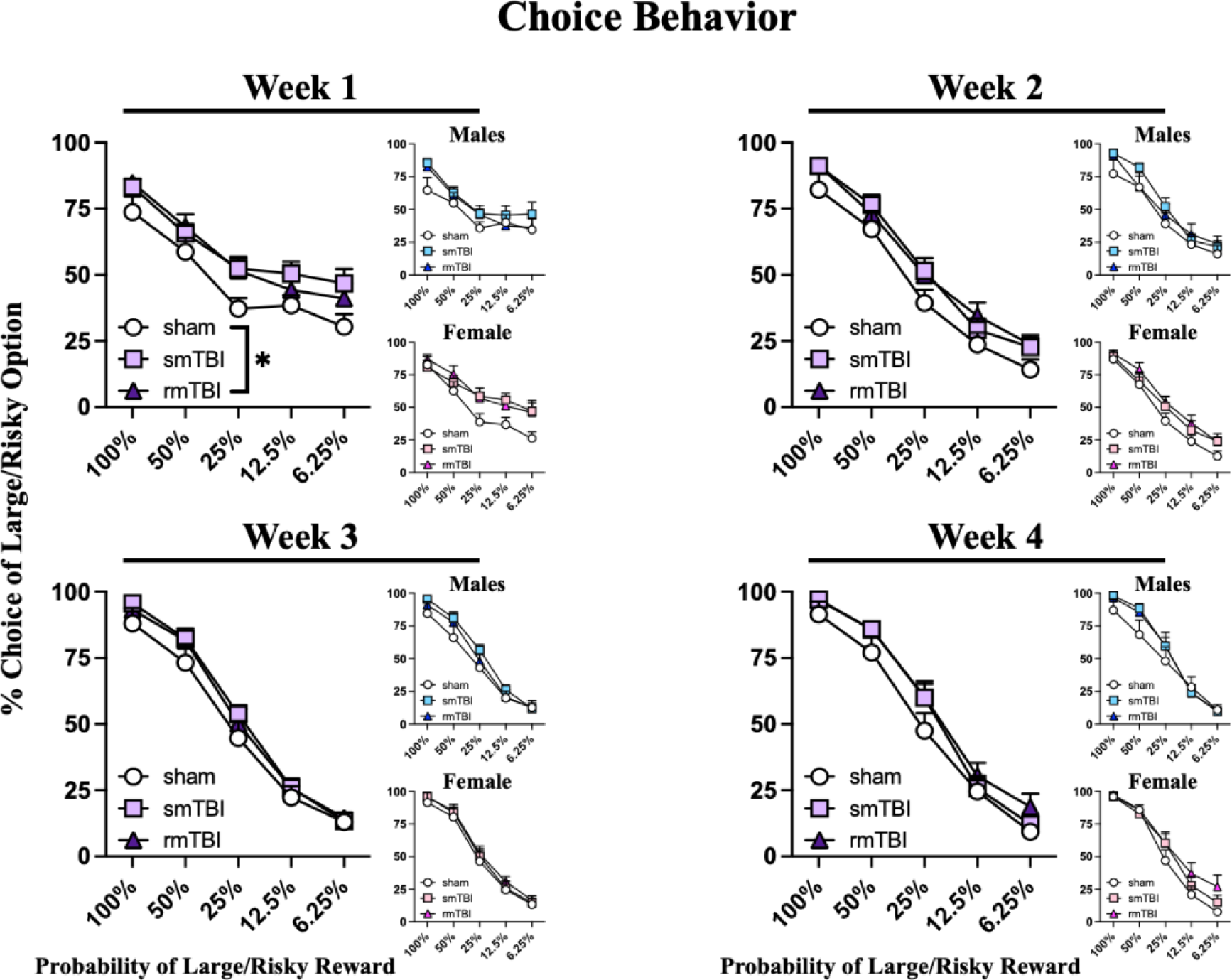
Choice behavior across four weeks post-final surgery. Line graphs represent percent choice of the large/risky option across five trial blocks. In week 1, a main effect of injury was observed when males and females were combined for analysis (purple graph). No differences in choice behavior were found between sham, single (smTBI), or repetitive (rmTBI) injury groups in weeks 2-4. Insets display choice behavior of males (blue) and females (pink) separately. No significant differences in choice behavior were found between sham, smTBI, or rmTBI groups for either males or females separately in weeks 1-4, although increased risky choice preference in injured females is clearly noted upon visual inspection within the first week post-final injury. Symbols represent mean ± SEM. * denotes p < 0.05 main effect of injury analyzed with three-way ANOVA.

In comparison to the effects observed in the first week following TBI, analysis of choice data from weeks 2-4 post injury failed to reveal a significant main effect of injury [F (2, 55) = 1.984, p = 0.147; F (2, 55) = 0.856, p = 0.431; F (2, 55) = 1.703, p = 0.192, respectively], sex [F (1, 55) = 0.170, p = 0.682; F (1, 55) = 0.952, p = 0.333; F (1, 55) = 0.667, p = 0.418, respectively] or injury x sex interaction [F (2, 55) = 0.213, p = 0.809; F (2, 55) = 0.254, p = 0.777; F (2, 55) = 0.177, p = 0.838, respectively]. Likewise, these analyses failed any other interactions of block x injury, block x sex or block x injury x sex (p > 0.05) in weeks 2-4. Collectively, these data indicate that TBI increases risky choice during probabilistic discounting, but these effects dissipate after extended re-training.

### Effects of mTBI on Choice Latency

Separate analyses compared how TBI affected choice latencies. Across weeks 1-4 of testing (Fig. 4), analysis of choice latency revealed a significant main effect of sex [F (1, 55) = 4.008, p = 0.050; F (1, 55) = 9.535, p = 0.003; F (1, 55) = 8.993, p = 0.004; F (1, 55) = 8.561, p = 0.005, respectively], reflecting that females were generally slower to make choices compared to males. However, these analyses did not yield a main effect of injury [F (2, 55) = 0.088, p = 0.915; F (2, 55) = 0.413, p = 0.664; F (2, 55) = 0.861, p = 0.428; F (2, 55) = 0.888, p = 0.417, respectively] or injury x sex interaction [F (2, 55) = 1.595, p = 0.212; F (2, 55) = 1.516, p = 0.229; F (2, 55) = 0.830, p = 0.441; F (2, 55) = 0.245, p = 0.783, respectively]. There were no other interactions of block x injury, block x sex, and block x injury x sex (p > 0.05) in weeks 1-4.

**Figure 4.**
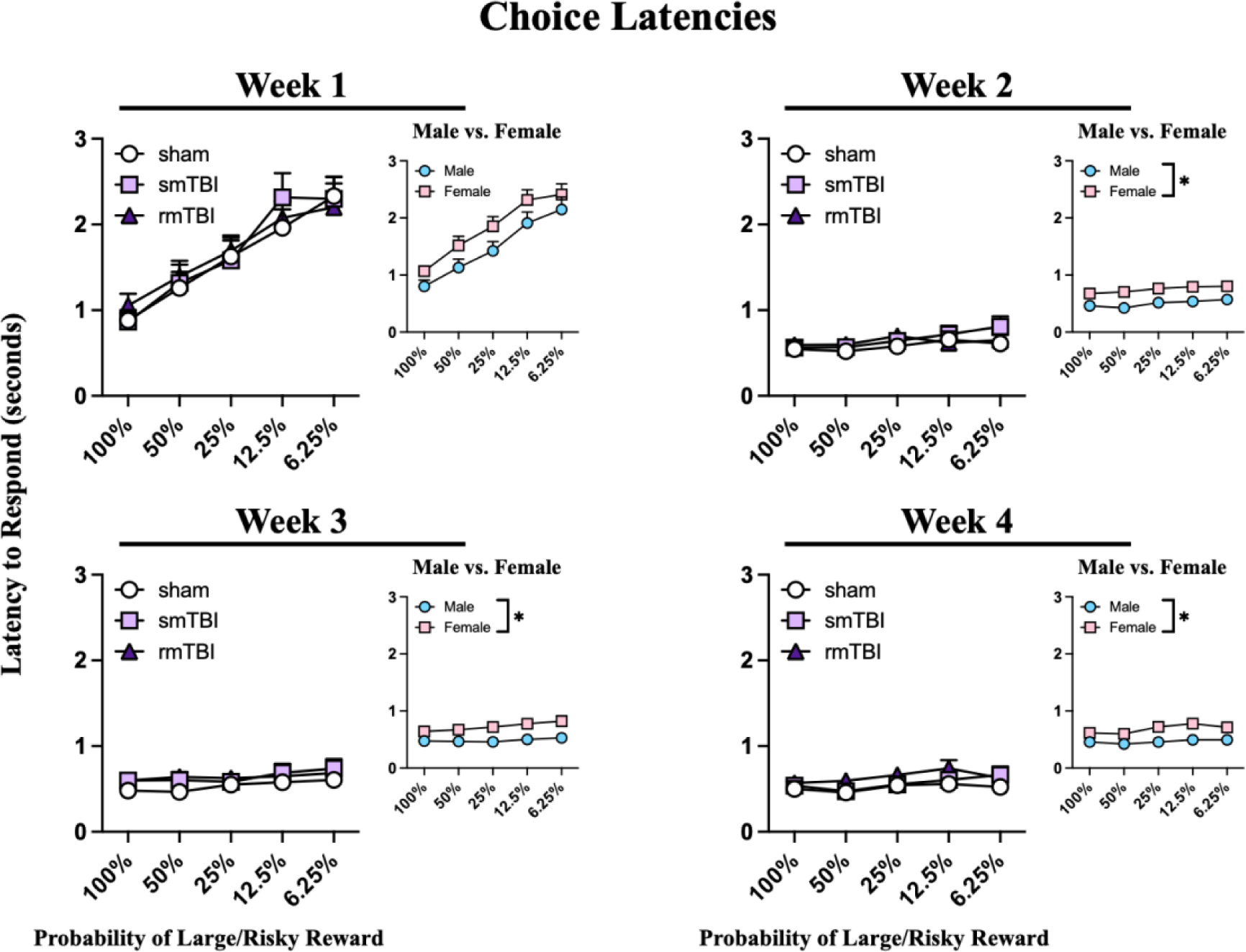
Choice latencies across four weeks post-final surgery. Line graphs represent choice response latencies across five trial blocks. No differences in choice latencies were found between sham, single (smTBI), or repetitive (rmTBI) injury groups in weeks 1-4 post-surgery. Insets display overall male vs. female choice latencies across all trial blocks. Across all 4 weeks, females demonstrated slower choice latencies compared to males. Symbols represent mean ± SEM. * denotes p < 0.05.

Since we found clear differences in choice latencies between males and females, we wanted to see whether there were differences in latencies to choose different options. Subsequent three-way ANOVA analyses were therefore conducted to compare choice latencies of trials that resulted in a risky choice versus a certain one (Fig. 5). Across weeks 1-4 of testing, analysis of choice latency revealed a significant main effect of sex [F (1, 55) = 4.600, p = 0.0364; F (1, 55) = 7.752, p = 0.0073; F (1, 55) = 5.119, p = 0.0276; F (1, 55) = 8.500, p = 0.0051, respectively], reflecting, again, that females were slower to make choices compared to males. However, these analyses did not yield a main effect of choice type [F (1, 55) = 0.6358, p = 0.4287; F (1, 55) = 1.388, p = 0.2438; F (1, 54) = 2.152, p = 0.1482; F (1, 55) = 3.731, p = 0.0586, respectively], injury [F (2, 55) = 0.2468, p = 0.7822; F (2, 55) = 0.1290, p = 0.8792; F (2, 55) = 1.391, p = 0.2576; F (2, 55) = 1.409, p = 0.2531, respectively] or injury x sex interaction [F (2, 55) = 1.303, p = 0.2800; F (2, 55) = 1.641, p = 0.2036; F (2, 55) = 0.9807, p = 0.3815; F (2, 55) = 0.9509, p = 0.3926, respectively]. There were no other interactions of choice x injury, choice x sex, and choice x injury x sex (p > 0.05) in weeks 1-4.

**Figure 5.**
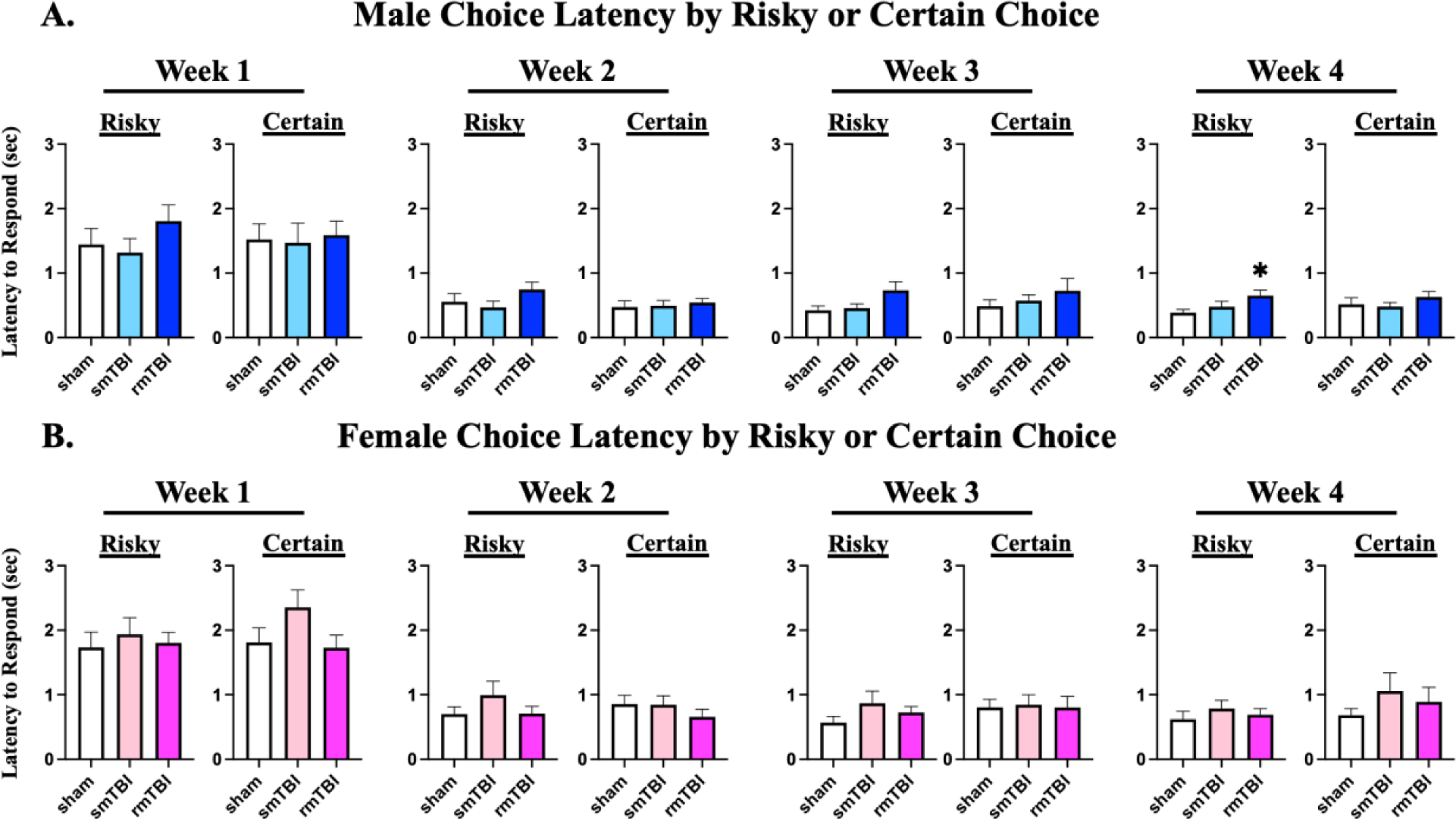
Choice latencies separated by choice type across four weeks post-final surgery. Bar graphs represent the averaged total percentage of choice latencies for trials that ended in either a risky or certain choice across all trial blocks. **A)** Males: No differences in choice latencies were found between sham, single (smTBI), or repetitive (rmTBI) injury groups in weeks 1-2 post-final surgery. In week 3, rmTBI tended to increase choice latency in risky choice trials, and this effect reached significance in week 4. **B)** Females: No differences in choice latencies were found between sham, smTBI, or rmTBI injury groups in in weeks 1-4. Bars represent mean ± SEM. * denotes p < 0.05 from shams analyzed with Dunnet’s multiple comparisons tests.

When the latencies of risky and certain choices were assessed separately, males, regardless of injury condition, showed no differences in choice latencies of trials that ended in a risky [F (2, 28) = 1.091, p = 0.3497; F (2, 28) = 1.495, p = 0.2417, respectively) or certain [F (2, 28) = 0.05336, p = 0.9481; F (2,28) = 0.1916, p = 0.8267, respectively] choice in weeks 1 and 2 post-final injury (Fig. 5A). However, in week 3, there was a strong trend towards increased latencies in trials that ended in risky choices (F (2, 27) = 3.194, p = 0.0569), and by week 4, this effect of mTBI was significant (F (2, 28) = 3.368, p = 0.0489). Dunnett’s multiple comparisons analysis revealed that rmTBI rats were slower to make riskier choices (p = 0.0297) compared to sham animals. No significant differences in choice latencies between male injury groups were observed during trials that ended in certain (p > 0.1) choices in weeks 3 and 4. In comparison, there were no significant differences between female injury groups in choice latencies of trials that ended in either a risky or certain (all ps > 0.1) choice across weeks 1-4 (Fig. 5B).

### Effects of mTBI on TH and NET

In a separate group of rats that did not undergo behavioral testing, we examined the effects of TBI on expression of markers associated with catecholamine transmission (Fig. 6). Analysis of protein levels within the mPFC and ACC collected from tissue 48 hours after final surgery revealed no significant changes in TH levels in males [F (2, 18) = 0.170, p = 0.8454; F (2, 17) = 0.440, p = 0.6512, respectively] or females [F (2, 21) = 0.415, p = 0.6658; F (2, 17) = 0.799, p = 0.4661, respectively] following injury. In the OFC, there were no significant changes in TH [F (2, 20) = 0.945, p = 0.4054] levels in males. However, in the OFC of females, we did observe a significant effect of injury on TH levels [F (2, 20) = 3.593, p = 0.0464]. Dunnett’s multiple comparisons analysis determined that smTBI rats displayed a significant increase in TH levels (p = 0.0296) compared to sham rats. Although rmTBI did not cause a statistically significant increase in female OFC TH levels, it is notable that these treatments did alter these levels in the same direction as smTBI, causing an 89.5% increase relative to shams.

**Figure 6.**
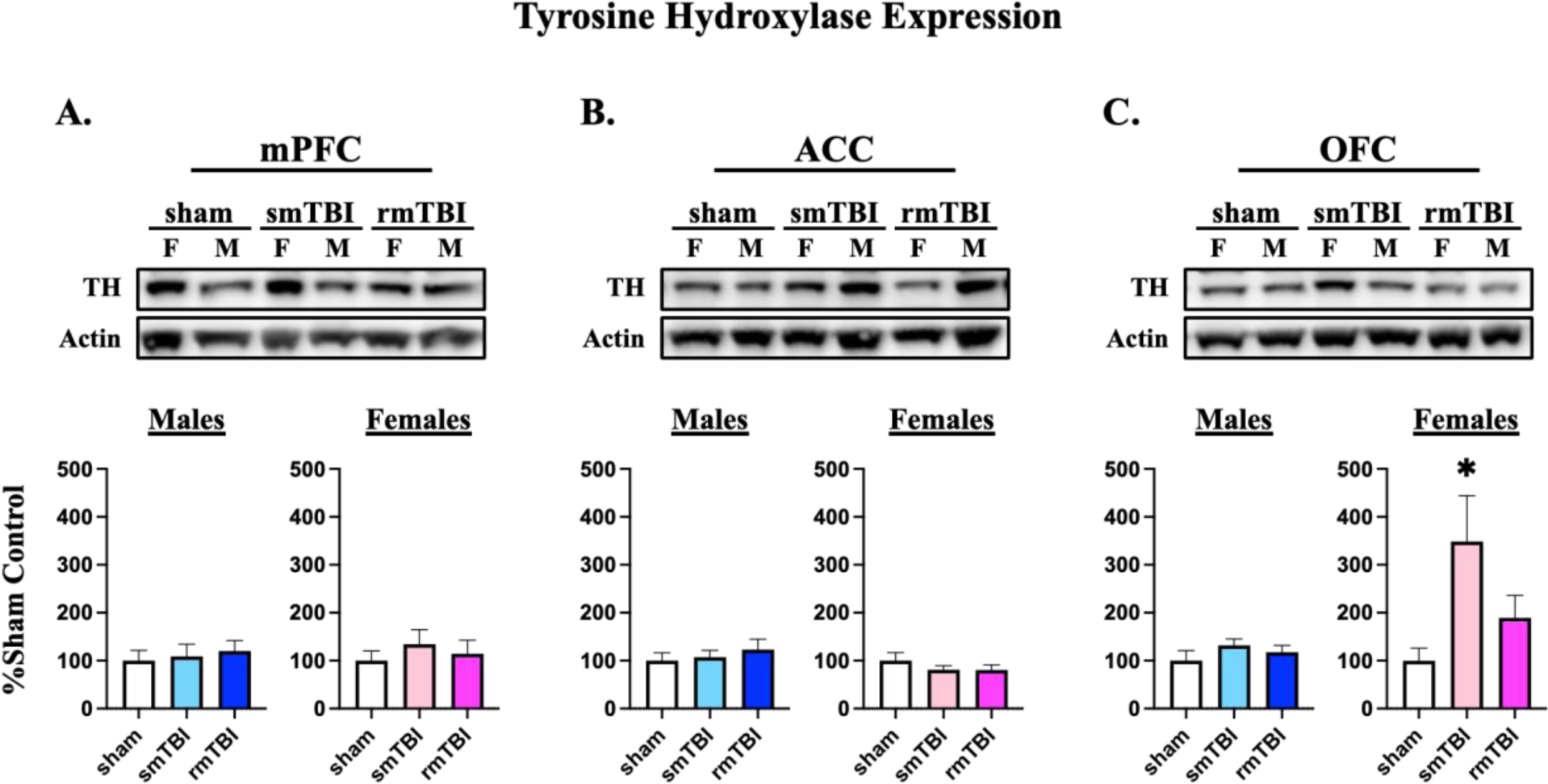
Male and female levels of tyrosine hydroxylase (TH) in the **A)** medial prefrontal cortex (mPFC), **B)** anterior cingulate cortex (ACC), and **C)** orbitofrontal cortex (OFC). Graphs represent percent total protein levels at 48 hours post-final surgery. No differences in TH levels were found between sham, single (smTBI), or repetitive (rmTBI) injury groups in the mPFC or ACC of either sex. In the OFC, smTBI significantly increased TH levels in females only. Bars represent mean ± SEM. * denotes p < 0.05 from sham analyzed with Dunnet’s Multiple Comparisons Tests.

Analysis of NET protein levels (Fig. 7) within the mPFC and ACC revealed TBI caused no significant changes in NET levels in males [F (2, 19) = 1.166, p = 0.3328; F (2,19) = 0.368, p = 0.6971, respectively] or females [F (2,19) = 0.558, p = 0.5814; F (2,21) = 0.545, p = 0.5879, respectively]. On the other hand, in the OFC, there was a significant effect of injury on NET levels that was apparent in both males [F (2, 12) = 3.953, p = 0.0480] and females [F (2, 17) = 3.854, p = 0.0417]. Dunnett’s multiple comparisons determined that rmTBI induced a significant decrease in NET (p < 0.05) levels compared to sham rats.

**Figure 7.**
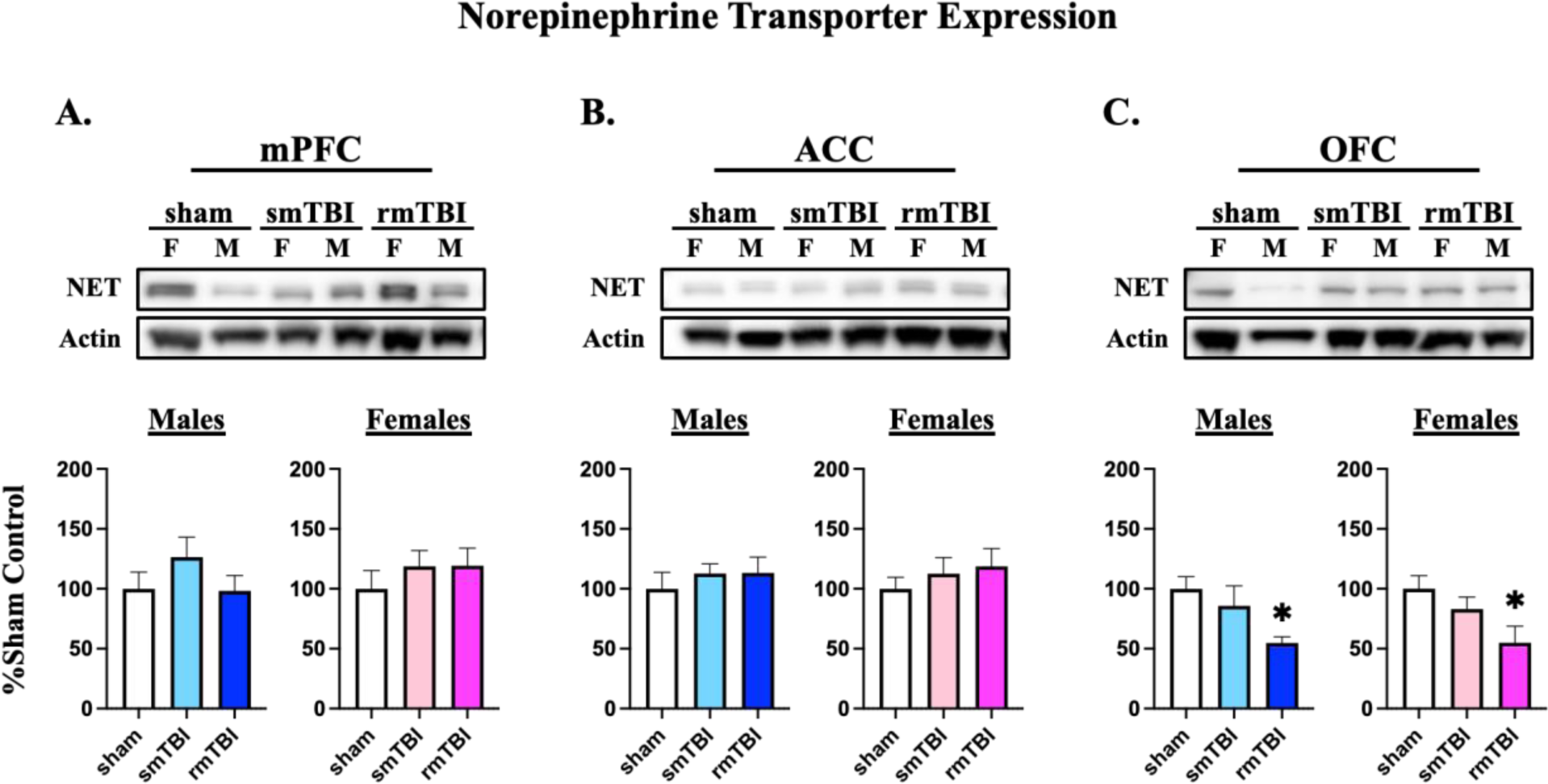
Male and female levels of norepinephrine transporter (NET) in the **A)** medial prefrontal cortex (mPFC), **B)** anterior cingulate cortex (ACC), and **C)** orbitofrontal cortex (OFC). Graphs represent percent total protein levels at 48 hours post-final surgery. No differences in NET levels were found between sham, single (smTBI), or repetitive (rmTBI) injury groups in the mPFC or ACC of either sex. In the OFC, rmTBI significantly reduced NET levels in both males and females. Bars represent mean ± SEM. * denotes p < 0.05 from sham analyzed with Dunnet’s Multiple Comparisons Tests.

## Discussion

Using a rodent assay of probabilistic discounting, the present study evaluated the effects of rmTBI on risk/reward decision making. During the first week following injury, mild TBI increased risky choice, and this effect achieved statistical significance in smTBI animals. Upon further inspection, it appeared that these increases were more pronounced in the female injury groups. Additionally, with extended testing, these effects resolved by week 2 post-injury indicating that the effects of mTBI on choice behavior are transient. While the effects of single injury align with previous TBI research, this study offers new insights into the effects of repetitive injury on risk/reward decision making. rmTBI does in fact disrupt adjustments in choice biases in response to changes in the relative risk of not obtaining rewards, but to a slightly lesser degree than a single impact.

Choice behavior on this assay is guided in part by the mPFC, which exhibits robust neural activity during periods of anticipation and subsequent interpretation of risk-related outcomes^92–94^. Following decisions, the mPFC updates value representations based on changes in reward probabilities to facilitate more efficient choices^55,56,87^. Inactivation of the PFC increases risky choices on the PDT^55^ when reward probabilities decrease over a session, and also increases risk taking on other tasks^95,96^. The medial OFC (mOFC) has also been shown to exert influence over choice behavior through assessing reward values^48,97–101^ and mitigating the impact that large/probabilistic rewards exert on subsequent decisions^45,57^. Similar to the mPFC, inactivation of the mOFC in rodents leads to increased risky choice in the PDT^57^. During a probabilistic reversal learning task, disruption of mOFC activity impairs the ability to incorporate information from previous outcomes to guide subsequent actions^102^. TBI patients with both mPFC and mOFC damage demonstrate difficulty learning from previous mistakes, insensitivity to future consequence of risk-related choices, and overall increased risky behavior^18–20^. Based on these previous observations, it is likely that our findings of increased risky choice induced by mTBI are related to disruptions in mPFC and mOFC operations.

When choice latencies were examined, we found that males, were generally quicker to make choices than females in the PDT regardless of injury condition. A more detailed analysis of choice latencies partitioned by whether rats made risky or certain choices, revealed a delayed effect of mTBI in males that approached statistical significance at week 3, and was fully apparent by week 4 post-final injury. Here, males that received repetitive injury showed increased hesitation to make risky choices. A previous study found that inactivation of the lateral OFC (lOFC) in males led to increased choice latencies in the PDT^55^, suggesting that these effects of rmTBI in males may be lOFC-related. Notably, patients with OFC damage have also demonstrated longer deliberation times in risk/reward decision making tasks^22,23^. These longer response times have been attributed to TBI patients having difficulty resolving competing options associated with uncertain or probabilistic outcomes^23,103^. As a result, these patients tend to make more disadvantageous choices when compared to non-injured individuals. Although these studies have focused primarily on single TBI cases, long-term deficits in information processing speed have been documented following repetitive TBIs^5,14^. Our results are consistent with these studies; however, our data indicates that these effects on response times do not occur immediately after injury, but rather develop over time. This finding is significant given that clinical studies often struggle with determining the onset of individual TBI-induced effects. Currently, we can only report that these longer choice latencies appear to manifest by week 4 after injury. Further research will be necessary to determine the full duration of these effects.

Interestingly, females appeared to be relatively impervious to injury-induced changes in choice latencies. It is known that males and females process reward-related information differently and use different strategies when making probabilistic-based decisions^101,104,105^. These differences in information processing can be attributed to sex differences in OFC activity during risk/reward decision making. Neuroimaging of the OFC during performance on the IGT found that males exhibit greater lOFC activity during task performance compared to females, while females exhibit mOFC-related activity during risk/reward decision making^106^. This would suggest that damage to the OFC may have a greater impact on lOFC-related processes, such as processing speed, in males than in females. Accordingly, one study that examined longitudinal patterns of decision making following pediatric TBI reported that injured males had difficulty processing information necessary to determine whether risks should be pursued, resulting in disadvantageous decisions, whereas females were still capable of making appropriate risk estimations^107^. Although these observations may explain why we did not observe changes in female choice latencies as a result of rmTBI, it is also possible that we are experiencing a ceiling effect with our female groups, making it difficult to detect potential changes in choice latencies. Future investigations are therefore needed to fully understand the sex-specific mechanisms underlying these differences following rmTBI.

The present study further investigated the effects of rmTBI on TH and NET protein levels within prefrontal regions that mediate risk/reward decision making processes. In the OFC, smTBI produced dramatic increases in TH within female, but not male, rats. While not statistically significant, rmTBI produced a roughly 90% increase in TH expression in females only. rmTBI also significantly reduced levels of NET in the OFC of both males and females. These results suggest that the OFC may be more susceptible to catecholamine dysregulation following repetitive mild head injuries.

Previous studies have demonstrated increased TH and decreased DAT within the PFC and striatum following experimental TBI^74,75,82^. During impact, mechanical forces result in neuronal hyperexcitability and the opening of calcium channels, which enhances the activity of TH^108,109^. This activation of TH accelerates catecholamine synthesis, leading to the accumulation of large amounts of DA and NE within the tissue. This “catecholamine storm” is typically short-lived, often occurring immediately after injury and lasting for up to twenty-four hours in some brain regions^72,73,110,111^. In areas such as the striatum, this dramatic increase is followed by a hypo-dopaminergic state, which has been observed as long as two weeks following experimental TBI^74^. In comparison, TBI induces a different pattern of alterations in post-injury catecholamine levels within the PFC. Increased TH activity and catecholamines levels have been observed in the mPFC of male rats up to two weeks following TBI^71^. At four weeks, the same study noted that TH activity within the mPFC was reduced, suggesting that the PFC might not follow the traditional pattern of a brief catecholamine storm followed by a lasting depression of catecholamine synthesis. In response to this sustained increase in DA and NE synthesis, downregulation of transporter expression may serve as a compensatory mechanism to prevent overaccumulation of intracellular catecholamines. Increased cytosolic DA, in particular, can result in oxidative stress, neurotoxicity, and eventual death of catecholaminergic cells^112–114^.

Interestingly, rmTBI’s effects on TH were less robust than those of smTBI. Catecholaminergic neurons exposed to repetitive injury may not have sufficient stores of DA and NE by the third impact to mount another storm of the same magnitude produced by the first injury. Nevertheless, this increased expression of TH at 48 hours post-injury suggests that the PFC continues to experience an elevated catecholaminergic state following rmTBI, leading to a decrease in NET as a compensatory response to increased intracellular levels of DA and NE. This may also explain why there is a lack of change in NET levels following single injury. smTBI rats experienced only 48 hours of catecholaminergic disruption before regulatory protein levels were assessed, whereas rmTBI rats experienced eight days of irregular catecholamine activity from the first injury to 48 hours post the final impact. Therefore, reduced NET may be both a time-dependent response to increased TH as well as a rmTBI-driven response to maintain catecholamine homeostasis following prolonged exposure to catecholamine instability. The lack of changes in male TH expression at 48 hours post mTBI could suggest that males experience a shorter duration of elevated TH that subsides prior to our selected timepoint. Following rmTBI, it is possible that while levels of TH are no longer elevated 48 hours post-injury, the TH-induced increases of DA and NE may still linger resulting in the observed decreases in NET expression in male rats.

The decrease in OFC NET expression induced by TBI would be expected to result in enhanced NE levels within this region. With this in mind, it is interesting to note that enhancing NE transmission with systemic treatment of the α2 antagonist yohimbine increased risky choice in a manner comparable to the effects reported here, using a version of the PDT similar to that used in this study (Montes et al., 2015). Although the specific neural locus where pharmacological enhancements in NE activity may promote risky choice remains to be clarified, the similarity of effects observed this previous study, and the current results allude to the possibility that abnormal increases in OFC NE transmission may contribute to alterations in risk/reward decision making induced by TBI.

The lack of protein changes in the mPFC and ACC were initially surprising, given that TBI-induced alterations in catecholaminergic activity have been reported in the mPFC^71^. One explanation could be that our current injury parameters may be sub-threshold for altering catecholamine regulation in these regions mPFC and ACC. The previously mentioned study of increased TH activity and levels of catecholamines in the mPFC was performed using an open skull CCI model^71^. Open-skull models of TBI often induce a more severe injury than a closed skull which can lead to increased detection of TBI-induced effects; however, these models are less suitable for repetitive TBI studies. Increasing our injury parameters to induce a stronger form of rmTBI may allow us to detect changes in TH and NET expression within these sub-regions. Nevertheless, this injury model has proved effective in revealing the susceptibility of the OFC to catecholamine dysregulation following TBI as well as the impact of rmTBI on PFC-related cognitive processes. Future studies will be designed to further evaluate these behavioral and neurochemical consequences, including the direct measurement of DA and NE levels at 48 hours post single and repetitive TBI.

## Conclusion

Overall, we report that mTBI produces transient increases in risk preferences, which appear to be largely driven by our female injury groups. While the effects of rmTBI are not as robust as those of a single impact, repetitive injury does appear to disrupt adjustments in choice biases in response to changes in the value of large/risky rewards. Additionally, we found that males are more likely to experience delayed disruptions in cost/benefit evaluations, resulting in longer deliberation periods when making risky choices. In practical terms, these pre-clinical findings suggest that individuals experiencing repetitive concussions should be made aware of tendencies toward risky choices following injury. In head injured males, longer deliberation times could by themselves be debilitating when faced with cost/benefit decisions. By evaluating specific subregions of the PFC, we determined that the OFC is more susceptible to catecholamine instability after rmTBI, a finding indicating that not all areas of the PFC contribute equally to the observed TBI-induced catecholamine imbalances. These time-dependent and sex-specific changes in risk/reward decision making and catecholamine regulation underscores the importance of evaluating both males and females in pre-clinical TBI studies. This work also demonstrates the effectiveness of combining this repetitive injury model with an operant-based behavioral paradigm to produce differential effects in male and female subjects. Future research with this model will allow for further investigation into the underlying mechanisms responsible for these behavioral changes as well as for testing potential treatment strategies to alleviate rmTBI-induced cognitive deficits.

## Transparency, Rigor, and Reproducibility Statement

All experimental procedures were in accordance with the Rowan University School of Osteopathic Medicine Institutional Animal Care and Use Committee and the National Institutes of Health Guide for the Care and Use of Laboratory Animals. For behavioral experiments, sample size was 9-11 animals per group (male: n = 11 sham, n = 10 smTBI, n = 10 rmTBI; female: n = 11 sham, n = 9 smTBI, n = 10 rmTBI). All training and testing procedures involving the PDT were adapted and verified based on studies previously described^55–57,83–87^. When assigning animals to either sham, smTBI, or rmTBI groups, a reverse Latin square method was used to limit bias by having equal amounts of risky and risk-adverse rats in each group. For Western blotting experiments, sample size was 8 animals per group (male: n = 8 sham, n = 8 smTBI, n = 8 rmTBI; female: n = 8 sham, n = 8 smTBI, n = 8 rmTBI). All reagents and antibodies used in this study are commercially available. Data was excluded only if there were bubbles/imperfections in the band that interfered with signal detection. Therefore, the “n” for all Western blotting data might vary from the total number of animals in each injury group but can be determined by the degrees of freedom provided in the Results section. These exclusion criteria are consistent with previous studies^89–91^. Normal distribution of primary behavioral and Western data was confirmed using Shapiro-Wilk tests.

## Acknowledgements

The authors thank Douglas Fox and Anuradha Krishnan for their technical assistance in this study. Figure 1 schematic was created using BioRender.com (accessed on 12/07/2023).

## Authors’ Contributions

Conceptualization: C.P.K. and R.L.N. Data Curation: C.P.K. and R.L.N. Formal analysis: C.P.K. Investigation: C.P.K and E.P. Methodology: C.P.K., J.A.L., R.R., S.B.F, and R.L.N. Project administration: C.P.K. and R.L.N. Software: C.P.K., S.B.F., and R.L.N. Supervision: R.L.N. Validation: C.P.K. and R.L.N. Visualization: C.P.K. Writing – original draft: C.P.K. Writing – review & editing: C.P.K., E.P., J.A.L., R.R., S.B.F., B.D.W., and R.L.N.

